# Leveraging a synthetic biology approach to enhance BCG-mediated expansion of Vγ9Vδ2 T cells

**DOI:** 10.1101/2025.05.05.651767

**Authors:** Christine M. Qabar, Lucas Waldburger, Jay D. Keasling, Dan A. Portnoy, Jeffery S. Cox

**Affiliations:** Department of Plant and Microbial Biology, University of California Berkeley, Berkeley CA 94720, USA; Department of Bioengineering, University of California Berkeley, Berkeley CA 94720, USA; Department of Electrical Engineering and Computer Science, University of California Berkeley, Berkeley CA 94720, USA; Department of Chemical and Biomolecular Engineering, University of California Berkeley, Berkeley CA 94720, USA; Joint BioEnergy Institute, Lawrence Berkeley National Laboratory, Emeryville CA 94608, USA; Biological Systems and Engineering Division, Lawrence Berkeley National Laboratory, Berkeley, CA, 94720, USA; Department of Molecular and Cellular Biology, University of California Berkeley, Berkeley CA 94720, USA

## Abstract

There is an urgent need to develop a more efficacious anti-tuberculosis vaccine as the current live-attenuated vaccine strain BCG fails to prevent pulmonary infection in adults. Our long-term goal is to test whether increasing the immunogenicity of BCG will improve vaccine effectiveness while maintaining its proven safety profile. In this study, we leverage a synthetic biology approach to engineer BCG to produce more (E)-4-hydroxy-3-methyl-but-2-enyl pyrophosphate (HMBPP), a phosphoantigen produced as an intermediate of bacterial—but not host—isoprenoid biosynthesis via the methylerythritol phosphate (MEP) pathway. Importantly, HMBPP strongly activates and expands Vγ9Vδ2 T cells, which are unique to higher-order primates and protect against *Mycobacterium tuberculosis* infection. Prior work to engineer BCG to produce specific ligands and antigens has been attempted to some success; however, our strategy exploits a self-nonself recognition mechanism in the host via HMBPP sensing, which has not been attempted before in this way. To inform the design of our recombinant strains, we performed synteny analyses of >63 mycobacterial species, which revealed that isoprenoid biosynthetic genes are not found in gene clusters or operons across all the 356 surveyed genomes. This analysis also revealed pair biases of isoprenoid biosynthesis genes frequently found in close proximity. In our engineering attempts, we found that simply overexpressing the rate-limiting gene in the pathway was toxic to the bacterium. Thus, we generated synthetic loci with the goal of specifically overproducing HMBPP, and tested the ability of these engineered strains to induce human Vγ9Vδ2 expansion in an *in vitro* stimulation assay. We found that BCG expressing a rationally-designed, synthetic MEP locus did not enhance Vγ9Vδ2 T cell expansion over the wild-type vaccine strain, suggesting that ectopic expression of multiple MEP genes may result in feedback inhibition of the pathway. However we found that overexpression of the HMBPP synthase GcpE alone potently induced Vγ9Vδ2 T cell expansion and did not result in downregulation of other pathway genes, presenting a successful strategy to accumulate HMBPP and overcome feedback inhibition in this pathway. While much remains to be done to ultimately develop a more efficacious vaccine, our data present a promising system to improve upon the BCG platform. To our knowledge, this is the first work to attempt reengineering of the MEP pathway in BCG to improve vaccine efficacy.

## Introduction

*Mycobacterium tuberculosis* (*Mtb*), the causative agent of tuberculosis disease, has been persistent throughout the course of human history. Most notably in the late 1800s to early 1900s, *Mtb* infected nearly the whole population of Europe and resulted in ∼25% mortality (1). Despite the discovery of the bacterium by Robert Koch over 142 years ago, *Mtb* remains a global health threat, killing approximately six thousand individuals per day and latently infecting an estimated quarter of the world’s population (2).

There is an urgent need to develop new vaccines with greater efficacy than the current live-attenuated vaccine strain Bacille Calmette-Guérin (BCG), an attenuated version of *Mycobacterium bovis* lacking several *Mtb*-specific virulence genes (3, 4). BCG has been used globally since 1921 despite its failure to prevent pulmonary infection in adults (5, 6). Vaccines currently in the clinical pipeline harness myriad strategies, including live attenuated, inactivated, or recombinant BCG, as well as subunit, viral vector, and DNA vaccines (1). While these candidates show promise with six currently in phase III clinical trials, many challenges exist, both biological (correlates of protection, epitope selection, limited animal models) and epidemiological (ethics of human trials, exclusion of pregnant women) in nature (1).

Building on the existing BCG vaccine platform is appealing due to its established safety profile in humans, ease of production and transport, and relatively inexpensive cost at a median $0.24 per dose in 2023 (7). Other attempts to engineer BCG focus on heterologous expression of antigens and immune ligands, modification and encapsulation of surface molecules, and induction of bacterial lysis to varying degrees of protection (8–12). In contrast, our strategy is novel in that it exploits a specific metabolic sensing pathway in the host. Our approach is to enhance the immunogenicity of BCG by modulating the production of an endogenously-synthesized immunostimulatory molecule in the bacterium. Specifically, we focused on activating Vγ9Vδ2 T cells, a small but important subset of T cells that are unique to humans and non-human primates (13). Vγ9Vδ2 T cells constitute the most abundant subset of γδ T cells in humans, comprising 60-95% of all circulating γδ T cells and expanding to ∼50% of total blood T cells upon infection with pathogens including *Mtb* (14–16). These T cells respond specifically and potently to a unique diphosphate molecule called (E)-4-hydroxy-3-methyl-but-2-enyl pyrophosphate (HMBPP), which is produced by most bacteria as an intermediate of isoprenoid biosynthesis (17–21).

Isoprenoids represent a large, important class of biomolecules found in all domains of life. As many as 95,000 isoprenoid natural products have been identified to date including cholesterol, heme, and vitamin K (22, 23). In bacteria, isoprenoids support fundamental cellular processes, including membrane fluidity, cell wall synthesis, electron transport, stress response, and signaling and communication (24, 25). Isoprenoids are essential in *Mtb*, which relies on the isoprenoid decaprenyl phosphate for the synthesis of the essential cell wall components lipoarabinomannan and the mycolyl-arabinogalactan-peptidoglycan complex, as well as the “linker” molecules between arabinogalactan and peptidoglycan in the cell wall (26–28). Further, *Mtb* employs the virulence factors tuberculosinol and isotuberculosinol to arrest phagosomal maturation during infection (29–31). Together, isoprenoids are critical to all domains of life and represent essential molecules for *Mtb* growth and pathogenesis.

Despite wide structural and functional diversity, all isoprenoids arise from the fundamental precursor isopentenyl pyrophosphate (IPP), which can be synthesized via two independent pathways: the mevalonate (MEV) pathway, which is primarily found in eukaryotes, archaea, and some bacteria, or the methylerythritol phosphate (MEP) pathway, which is found in bacteria, plants, and algae (32, 33) (Figure 1A). Importantly, HMBPP is only produced via the MEP pathway, which is essential in *Mtb* (34).

**Figure 1:**
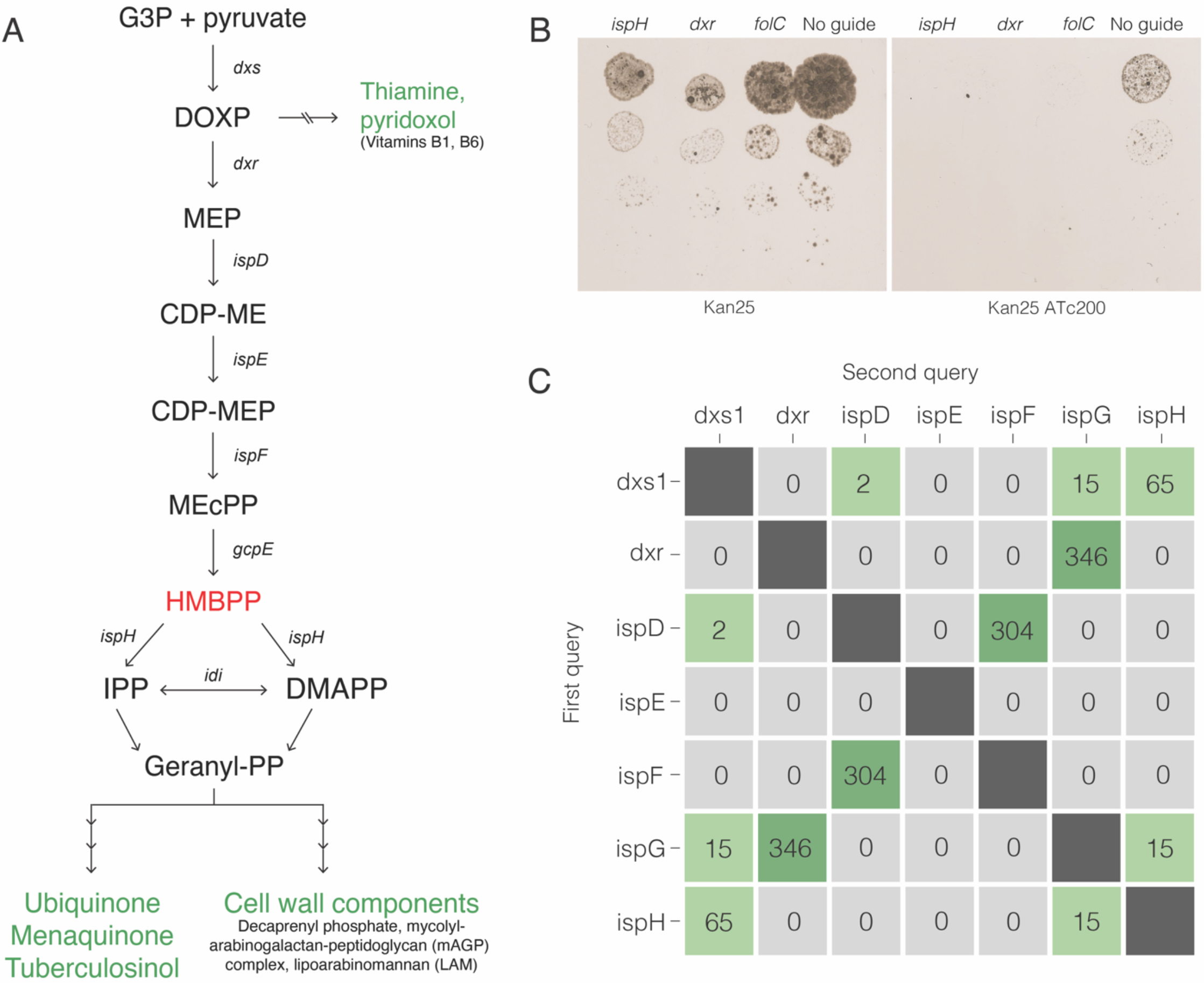
The MEP pathway is essential in BCG. **A. The MEP pathway of isoprenoid biosynthesis**. Shown in green are end products and shunts; shown in red is HMBPP, the activator of Vγ9Vδ2 T cells. Genes encoding enzymatic steps are as follows: *dxs*, DOXP synthase; *dxr*, DOXP reductoisomerase; *ispD*, MEP cytidylyltransferase; *ispE*, CDP-ME kinase; *ispF*, MEcPP synthase; *gcpE*, HMBPP synthase; *ispH*, HMBPP reductase; *idi*, isopentenyl diphosphate isomerase. **B. Silencing of MEP genes is lethal in BCG**. Targeting of *dxr* and *ispH* by inducible CRISPR interference was lethal in BCG. A sgRNA targeting the known essential gene *rpoB* and a non-targeting sgRNA were included as controls. sgRNAs were cloned into the vector pLJR965 carrying CRISPRi machinery and transformed into BCG. Transcriptional repression of target genes was achieved by the addition of ATc. Plates were incubated at 37°C for a minimum of 12 days before counting and imaging. **C. Synteny analysis reveals naturally-occurring gene pair biases to leverage in our synthetic platform**. Several gene pairs are frequently found in proximity to each other across mycobacteria: *dxs1 + ispH*; *dxr + ispG/gcpE*; *ispD + ispF*; and occasionally *ispH + ispG/gcpE*. Synteny analysis was performed on 353 mycobacterial genomes using each gene as a query to generate this 7x7 matrix of co-occurrence.

The Vγ9Vδ2 T cell receptor (TCR) indirectly senses HMBPP via an “inside-out” mechanism. Via a yet-undetermined mechanism, bacterial HMBPP accesses the host cell cytosol where it binds a positively-charged groove on an intracellular domain of the transmembrane butyrophilin BTN3A1 in an antigen presenting cell (35–39). Upon HMBPP binding, the surface-associated BTN3A1:BTN2A1 heterodimer undergoes a conformational change which is recognized by the Vγ9Vδ2 TCR (14, 38). These noncanonical T cells also respond to endogenous host IPP (40), however HMBPP is 10,000-fold more potent an activator than IPP (41), demonstrating a considerable bias toward detecting microbial-derived metabolites. Together, the strength and specificity of this interaction suggests that the Vγ9Vδ2-phosphoantigen sensing pathway is likely predominantly a self-nonself detection mechanism.

Once activated, Vγ9Vδ2 T cells mount a robust immune response, including the production of important antimicrobial factors like granulysin, granzyme A, perforin, tumor necrosis factor alpha (TNF-α), granulocyte-macrophage colony-stimulating factor (GM-CSF), and interferon gamma (IFNγ) (42–47). Importantly IFNγ in turn leads to the production of reactive nitrogen species, which can control bacterial growth *in vivo* (48). Further, HMBPP-stimulated Vγ9Vδ2 T cells support the canonical αβ T cell response to infection by producing IL-12, the major cytokine driving Th1 polarization of T cells which is crucial to a protective anti-*Mtb* response (44, 49). Finally, activated Vγ9Vδ2 T cells have prolific memory recall abilities, expanding 60-fold in HMBPP-immunized non-human primates (44) and a reported memory sustained up to seven months post exposure (50). Importantly, activation of Vγ9Vδ2 T cells is protective *in vivo*. Studies in non-human primates have demonstrated that treatment with an HMBPP analog, immunization with HMBPP-producing *Lm*, or adoptive transfer of *ex vivo* activated Vγ9Vδ2 T cells protects against *Mtb* infection (44, 45, 51). Thus, HMBPP-mediated activation and subsequent expansion of Vγ9Vδ2 T cells is an attractive target for improving the BCG vaccine.

In this study, we leveraged a synthetic biology approach to metabolically engineer BCG to produce excess HMBPP and thus, stimulate a more robust Vγ9Vδ2 T cells response. A similar approach of expressing a synthetic MEP operon has been previously used to generate a “microbial cell factory” for economically-relevant isoprenoid production in *Bacillus subtilis* (52). However, in this study we leverage this approach to improve the immunogenicity of the BCG vaccine. We tested our recombinant strain in primary cells and found that a single-gene overexpression platform was most effective at stimulating Vγ9Vδ2 T cells, compared to overexpression of a synthetic MEP locus. Together, these data provide critical *in vitro* validation of metabolically reengineered strains of BCG, and can be built upon to develop a more efficacious anti-*Mtb* vaccine.

## Methods

### Bacterial strains and culture

*Mycobacterium tuberculosis* variant *bovis* BCG Pasteur (ATCC 35734) was routinely grown in Middlebrook 7H9 liquid medium or 7H10 agar (Difco) supplemented with 10% OADC (oleic acid-albumin-dextrose-catalase) and 0.05% Tween80. *E. coli* strains DH5α or NEB® 10-beta were grown in LB broth or agar and used for propagation and cloning of plasmids. When required, the following antibiotics were used: kanamycin (25 µg/ml for mycobacteria, 50 µg/ml for *E. coli*); hygromycin (50 µg/ml for mycobacteria, 150 µg/ml for *E. coli*); zeocin (25 µg/ml for mycobacteria, 50 µg/ml for *E. coli*); anhydrotetracycline (200 ng/ml).

### Molecular cloning

Primers, oligos, and plasmids used in this study are listed in Supplementary Table 1. Standard electroporation protocols were used for the transformation of plasmids into *E. coli* and mycobacteria. 50uL of electrocompetent DH5α *E. coli* was combined with 5ul of ligation product or plasmid DNA and incubated on ice for 30 min, then heat shocked at 42°C for 30 sec, chilled on ice for 5 min, and recovered at 37°C in Luria broth for one hour. Cells were then plated on selective LB agar and incubated 37°C overnight. Electrocompetent mycobacteria were prepared by washing mid-log (OD 0.5-0.8) cultures four times in decreasing volumes of sterile 10% glycerol. The final resuspension concentrates cultures ∼20-25x, and electrocompetent cells can be used fresh or frozen at -80°C. 200ul of electrocompetent cells were combined with 5ul of plasmid DNA in a 0.2cm electroporation cuvette, and cells were electroporated with a single pulse at 2.5 kV with 25 μF capacitance and 1,000 Ω resistance. Transformants were recovered overnight at 37°C in 7H9 media. The following day, cells were plated on selective 7H10 agar containing the appropriate antibiotic. 3-6 clones were randomly selected and verified via PCR.

### CRISPRi

Gene silencing and gene deletion were performed using the site-specific transcriptional repression system CRISPRi as previously described (53). Briefly, guide oligos were annealed and ligated into the integrative, dCas9-containing pLJR965 vector and the subsequent plasmid was transformed into BCG. Inducible transcriptional repression was achieved by treatment with 200 ng/ml anhydrotetracycline (ATc).

### Synteny analysis

To identify MEP synteny across mycobacterial genomes, we used the program ‘core analysis of syntenic orthologs to prioritize natural product gene clusters’ (CORASON) (54) with some updates. Briefly, 353 *Mycobacterium* genomes representing at least 62 unique species were downloaded from the NCBI Genome database (55). A database of protein sequences was generated using DIAMOND v4.0.515 (56) and queried with the IspE protein sequence from *Mycobacterium bovis* BCG. Results were filtered to include 5 protein sequences upstream and downstream of the IspE match. We created a second DIAMOND database of protein sequences within the neighborhood of IspE and queried with the remaining MEP protein sequences from BCG. The number of MEP genes within the gene neighborhood of IspE across the *Mycobacterium* genomes were counted and are represented in Figure 1C. To query whether MEP genes are operonic in any bacterium, we downloaded 36,193 bacterial reference genomes from NCBI and searched for all MEP genes in close genomic proximity. A new DIAMOND database was generated from the reference genomes and queried for the IspE protein sequence as previously described. Gene neighborhoods were searched for matches to the remaining MEP pathway genes.

### Generation of synthetic constructs

#### (I) Expression of a synthetic MEP locus (pCQ88)

The pMV306.hyg backbone was digested with KpnI and XbaI to generate non-compatible ends. The genes *dxs1, dxr, ispD, ispE, ispF*, and *gcpE* were included; HMBPP reductase *ispH* was not included in an attempt to optimize HMBPP accumulation. The six genes were further subdivided into two-gene operons, driven by unique strong promoters and preceding individual terminators, to optimize expression of each gene (Figure 2A). To ensure all genes are expressed, genes were assembled into two-gene operons informed by the genomic pair biases identified in our bioinformatic analysis (Figure 1C).

**Figure 2:**
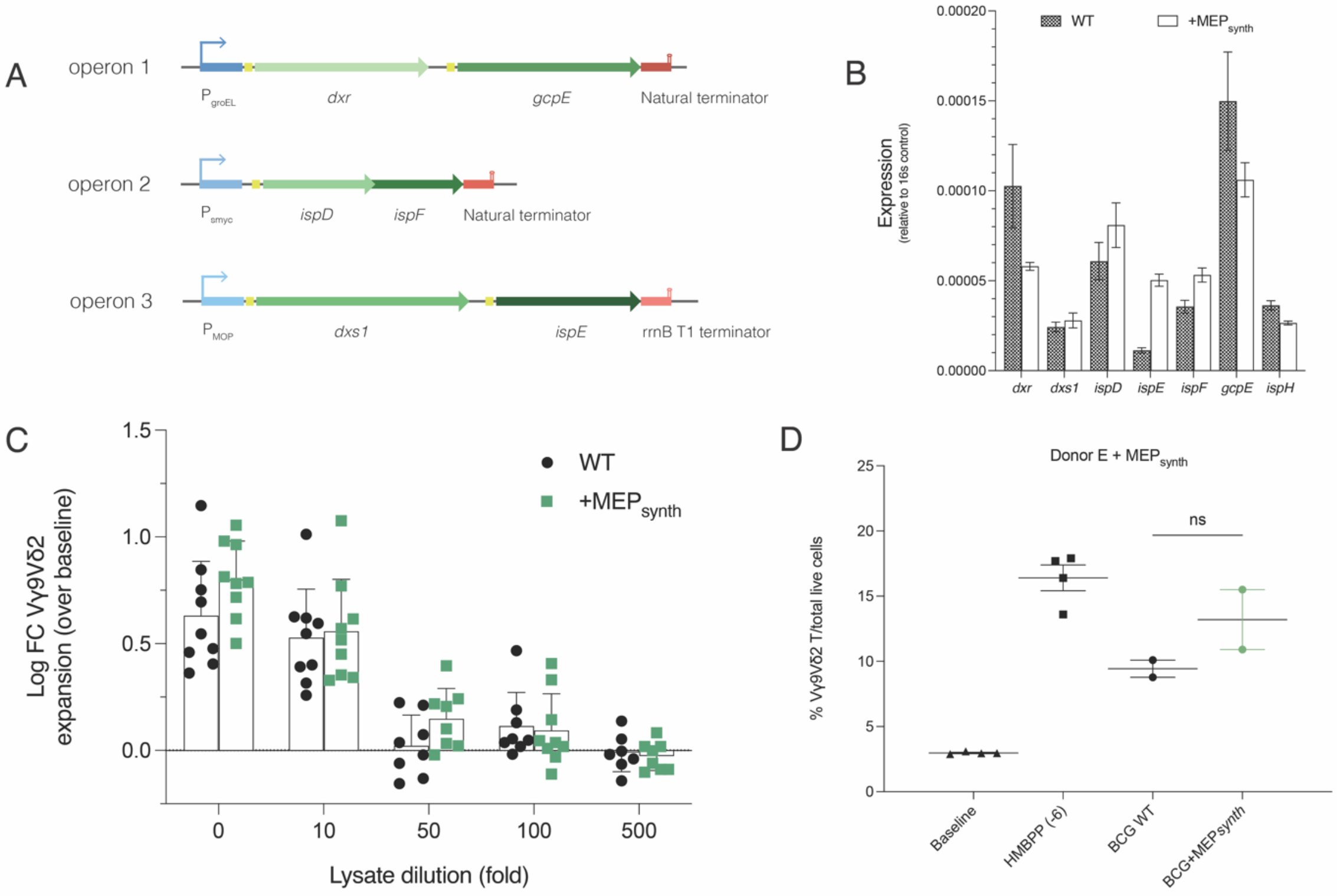
Generation of a synthetic MEP locus. **A. Designing pCQ88, a synthetic MEP locus**. Using gene pairs identified in our synteny analysis, we designed an integrative plasmid carrying an engineered biosynthetic gene cluster comprising all upper-branch MEP genes. Each operon is composed of one pair of genes (green) driven by a strong, unique mycobacterial promoter (blue), preceded by a ribosome binding site (yellow), and followed by either an exogenous or natural terminator (red). **B. Genes in the synthetic MEP locus are not strongly overexpressed**. No significant differences in individual gene expression exist between WT and BCG+pCQ88. RNA was isolated from WT and pCQ88-expressing BCG and used in a RT-PCR reaction to assess relative expression of each gene in the synthetic locus. Expression data for *ispH* is included as a control as it was not encoded in the synthetic MEP synthetic locus. N=3 biological replicates in technical triplicate. **C, D. The engineered biosynthetic gene cluster pCQ88 does not strongly enhance Vγ9Vδ2 T cell expansion over baseline**. There is no significant difference in Vγ9Vδ2 T cell expansion between WT and BCG+pCQ88. Shown is percent expansion of Vγ9Vδ2 T cells by pure HMBPP or lysate from either WT or BCG+pCQ88 at various dilutions **(C)** and at 1:500 diluted **(D)** compared to baseline. Data shown include data from three unique PBMC donors.

As *dxr* and *gcpE* are found within an operon in BCG, the genes were placed in an operon driven by PgroEL. Unique RBSs were placed upstream of each gene; the RBS from *groEL* upstream of dxr and the native RBS upstream of *gcpE*. 30 bp downstream of the *gcpE* stop codon were included to capture the native terminator. The genes *ispD* and *ispF* were placed in an operon as they are operonic in BCG, with overlapping coding regions. The strong mycobacterial promoter Psmyc and its associated RBS was placed upstream of ispD with no intergenic RBS due to the ORF overlap. 30bp downstream of the *ispF* stop codon were included to capture the native terminator. The genes *dxs1* and *ispE* were placed under control of the MOP promoter from pMH406. The Hops RBS was placed upstream of *dxs1* and the *gcpE* RBS was placed upstream of *ispE*. The rrnB T1 terminator from pMV261 was placed downstream of *ispE* to strongly terminate expression of the end of the three operons.

Resulting operons were optimized to achieve a GC content of <65% while preserving codon usage. Final, codon-optimized operons were ordered from Twist Bioscience (San Francisco, CA), amplified from the delivery vector, and cloned into a final vector via Gibson assembly. Transformants were confirmed via whole plasmid sequencing (Plasmidasaurus, San Francisco, CA).

#### (II) Overexpression of HMBPP synthase GcpE (pCQ105)

The pMV306.hyg backbone was digested as described above. The strong mycobacterial promoter Psmyc from the plasmid pUV15 was PCR amplified using primers oCQ404 and oCQ405 to introduce 5’ homology to the KpnI end of the digested vector and 3’ homology to the *gcpE* insert. The gene *gcpE* was PCR amplified from BCG gDNA using primers oCQ406 and oCQ407 to introduce 5’ homology to the Psmyc insert and 3’ homology to the XbaI end of the digested vector. All homologous overhangs are 20bp. The two inserts were assembled into the digested vector via Gibson assembly and transformed into NEB® 10-beta competent *E. coli*. Transformants were confirmed via whole plasmid sequencing.

#### RNA isolation and RT-PCR

Cultures were grown to mid-log (OD_600_ ∼1.0), pelleted, resuspended in 1 ml Trizol, and lysed via bead beating thrice in 30 second increments, incubating on ice for five minutes in between bursts. RNA was then extracted from the supernatant via chloroform-ethanol extraction, washed using a PureLink RNA Mini kit (Invitrogen), and treated with DNAse I. Following a 10 minute DNAse heat inactivation at 75°C, RNA was stored at -80°C. Primers were designed with the following criteria: 17-25 bp in length, 3’ G/C clamp, 75-150 bp amplicon, Tm ∼60ºC. Primers were also analyzed for self- and cross-secondary structure. cDNA was prepared using a SuperScript III First-Strand Synthesis Kit (Invitrogen). Briefly, random hexamer primers were annealed to template RNA, cDNA was synthesized using reverse transcriptase, and residual RNA was digested by RNAse H. cDNA was stored at -20°C. The SsoAdvanced Universal SYBR Green Supermix (BioRad) was used for RT-PCR reactions on genes of interest as well as 16S for normalization. cDNA samples were run in technical triplicate on a CFX Connect Real-Time System (BioRad) running CFX Manager software. Normalized expression ratios were obtained via the 2^-ΔΔCq^ (Livak) Method (57).

#### PBMC expansion assays

Expansion assays were performed as previously described (15, 34). Low-molecular-weight filtrates were generated by harvesting mid-log (OD_600_ ∼0.5) cultures, which were washed in DPBS and lysed via bead-beating in Lysing Matrix B tubes (MP Bio). Lysates were then pelleted and the supernatant fractionated using 3kDa filter cartridges (Amicon). Lysate supernatant fractions <3 kDa were used for subsequent assays as this fraction contains microbial HMBPP. Primary human peripheral blood mononuclear cells (PBMCs) were acquired from the Stanford Blood Center (Stanford, California, USA). Cells were cultured in R10 media (RPMI-1640 with 2mM glutamine, 10mM HEPES, 10% FBS, 50 U/ml pen-strep, and 50μM β-Mercaptoethanol) supplemented with 100 U/ml IL-2. PMBCs were plated at a density of 7.5x10^5^ cells/well in a U-bottom 96-well plate and incubated overnight at 37°C, 5% CO2, with humidity. The following day, PBMCs were stimulated with either pure HMBPP or bacterial lysates at various dilutions. Fresh media was added on day 4, and cells were harvested, stained with antibodies, and fixed on day 6. Flow cytometry was performed on day 7. Antibody panel:

| Marker | Fluorophore | Ratio | Clone | Source |
| --- | --- | --- | --- | --- |
| Vδ2 | PacBlue | 1:50 | B6 | Biolegend cat #331414 |
| Vγ9 | APC | 1:50 | B3 | Biolegend cat #331310 |
| CD3 | PerCP-Cy5.5 | 1:100 | OKT3 | Biolegend cat #317337 |
| $\alpha\beta$ TCR | FITC | 1:16.67 | IP26 | Biolegend cat #306706 |

## Results

### The MEP pathway is essential in BCG

Most Mycobacteria solely utilize the MEP pathway for isoprenoid biosynthesis, with the exception of a *M. marinum*-derived clade (58–60). Several genes in the MEP pathway are essential in *Mtb* (34, 61, 62), and because BCG and *Mtb* are closely related, we predicted that the MEP pathway is also essential in BCG. Indeed, we were unable to obtain a chromosomal deletion of *dxr* or *ispH*, two genes in the MEP pathway. To definitively demonstrate that these genes are essential, we turned to CRISPR interference (CRISPRi) to inducibly repress these genes in BCG. Using single guide RNAs that target *dxr* or *ispH*, as well as against the known essential gene *folC*, we found that repression of either gene inhibited growth (Figure 1A, B), supporting that the MEP pathway is essential in BCG. Further, transposon mutants of the MEP genes *ispH* and *dxs* in *Mtb* had impaired survival in a phagosome (63), underscoring the importance of the MEP pathway to viability and survival in a host. Together, these data represent the first evidence to demonstrate the essentiality of the MEP pathway in BCG, presenting barriers to future engineering of isoprenoid biosynthesis.

### Synteny analysis of MEP genes across mycobacterial genomes

Given the evidence supporting a protective role for Vγ9Vδ2 T cells in TB infection, we hypothesized that enhanced Vγ9Vδ2 activation might also enhance protection. To engineer a strain of BCG that produced elevated levels of HMBPP, we initially set out to combine all MEP genes leading to HMBPP synthesis into a single locus to maximize expression. To rationally design this synthetic MEP locus, we first sought to determine whether any bacterial species already encode all MEP genes either in proximity or in an operon. Most bacteria do not encode the MEP pathway as an operon (64), in agreement with our finding only one relevant hit: *Nostoc sphaeroides*, a non-pathogenic environmental microbe (data not shown). This suggests that in all bacteria—and more saliently, in all mycobacteria—the MEP genes are distributed across the genome. To determine whether any MEP genes are found together in mycobacteria, we queried 353 mycobacterial genomes for co-occurrences of MEP genes clustered in genome neighborhoods. The synteny analysis revealed several gene pairs that are frequently found in proximity: *dxs1 + ispH; dxr + ispG/gcpE; ispD + ispF*; and occasionally *ispH + ispG/gcpE* (Figure 1C). Because these pair biases are strongly represented across the genus, we sought to preserve them in our subsequent engineering attempts.

### Generation of a synthetic MEP locus

Our synteny analysis findings posed a unique challenge: if all MEP genes in all bacteria are scattered across the genome, we must carefully design a synthetic locus which takes into account the tendency for certain gene pairs to be found in proximity to one another, as well as the potential that each gene in the MEP pathway is regulated under vastly different conditions. Using the pair biases revealed in our synteny analysis, we designed the plasmid pCQ88, an integrative plasmid carrying three, two-gene operons composed of strong promoters, defined ribosome binding sites, and strong terminators to prevent readthrough transcription (Figure 2A). We transformed this plasmid expressing the six genes *dxr, gcpE, ispD, ispF, dxs1, ispE* into BCG and confirmed successful integration via junction PCR.

To determine if this strain was more stimulatory to Vγ9Vδ2 T cells, we turned to an *in vitro* expansion assay. Animal models were not at our disposal because Vγ9Vδ2 T cells are restricted to higher order primates (65). In our *in vitro* expansion assay, human peripheral blood mononuclear cells (PBMCs) were treated *ex vivo* with lysates from either WT BCG or a pCQ88-carrying strain and assessed for Vγ9Vδ2 expansion via flow cytometry. We found that BCG containing pCQ88, our first iteration of a synthetic MEP locus, modestly enhanced Vγ9Vδ2 expansion over baseline over WT (Figure 2C, D). This small boost in expansion over WT was not unexpected, as these genes are not natively encoded together and thus, our synthetic locus approach was ambitious. Indeed, upon qRT-PCR analysis, we observed that not all of the ectopically-expressed genes were in fact overexpressed over WT baseline. Notably, we observed downregulation of *dxr* and *gcpE*, two critical steps in the isoprenoid biosynthetic pathway (Figure 2B), which likely restricted the production of HMBPP and magnitude of subsequent Vγ9Vδ2 T cell expansion.

One limitation to our synthetic platform of MEP pathway expression is the potential for feedback inhibition by IPP, DMAPP, and other downstream metabolites. Intracellular levels of MEP intermediates are tightly regulated via negative feedback mechanisms (66, 67). Thus, it is possible that ectopic expression of the synthetic locus overwhelmed the cell with metabolites such that the cell throttled all IPP synthesis. Indeed we observed downregulation of *ispH*, which was not ectopically expressed in the synthetic locus, suggesting that even the native pathway may have been undergoing feedback inhibition (Figure 2B). This is in line with a mild growth defect of BCG carrying pCQ88 (data not shown). Overall, this synthetic MEP locus did not perform as well as expected, and thus we turned to a new strategy to improve upon it.

### Generation of a *gcpE* single gene overexpression platform

To avoid burdening the cell by overexpressing multiple metabolic genes, we turned to a simpler approach in which only one gene is ectopically expressed. We attempted to overexpress *dxr* as it is the first committed step in the pathway; however, our initial attempts to transform a *dxr*-overexpressing plasmid into BCG failed, whether *dxr* was under the control of a strong promoter or its native promoter. We were able to obtain transformants when *dxr* was placed under an anhydrotetracycline-inducible promoter, but observed no growth upon induction (data not shown). Thus it appears that overexpression of *dxr* alone is toxic in BCG, which has been reported previously in *Mtb* (34). This is likely because accumulation of early MEP intermediates is toxic. We instead expressed the HMBPP synthase GcpE to specifically accumulate HMBPP, as it likely will not accumulate other intermediates in the pathway (Figure 3A). A similar strategy was successfully engineered in *Mtb* (34), providing strong evidence that this approach would improve Vγ9Vδ2 T cell expansion in our system. In contrast to our synthetic MEP strain, this *gcpE* overexpression strain had no growth defect (data not shown). We confirmed overexpression of *gcpE* and found no global pathway repression, in contrast to our previous construct (Figure 3B). Interestingly, we observed overexpression of the kinase IspE in our engineered strain in both constructs (Figures 2B, 3B), suggesting that expression of this gene may be responsive to a yet-undetermined stimulus.

**Figure 3:**
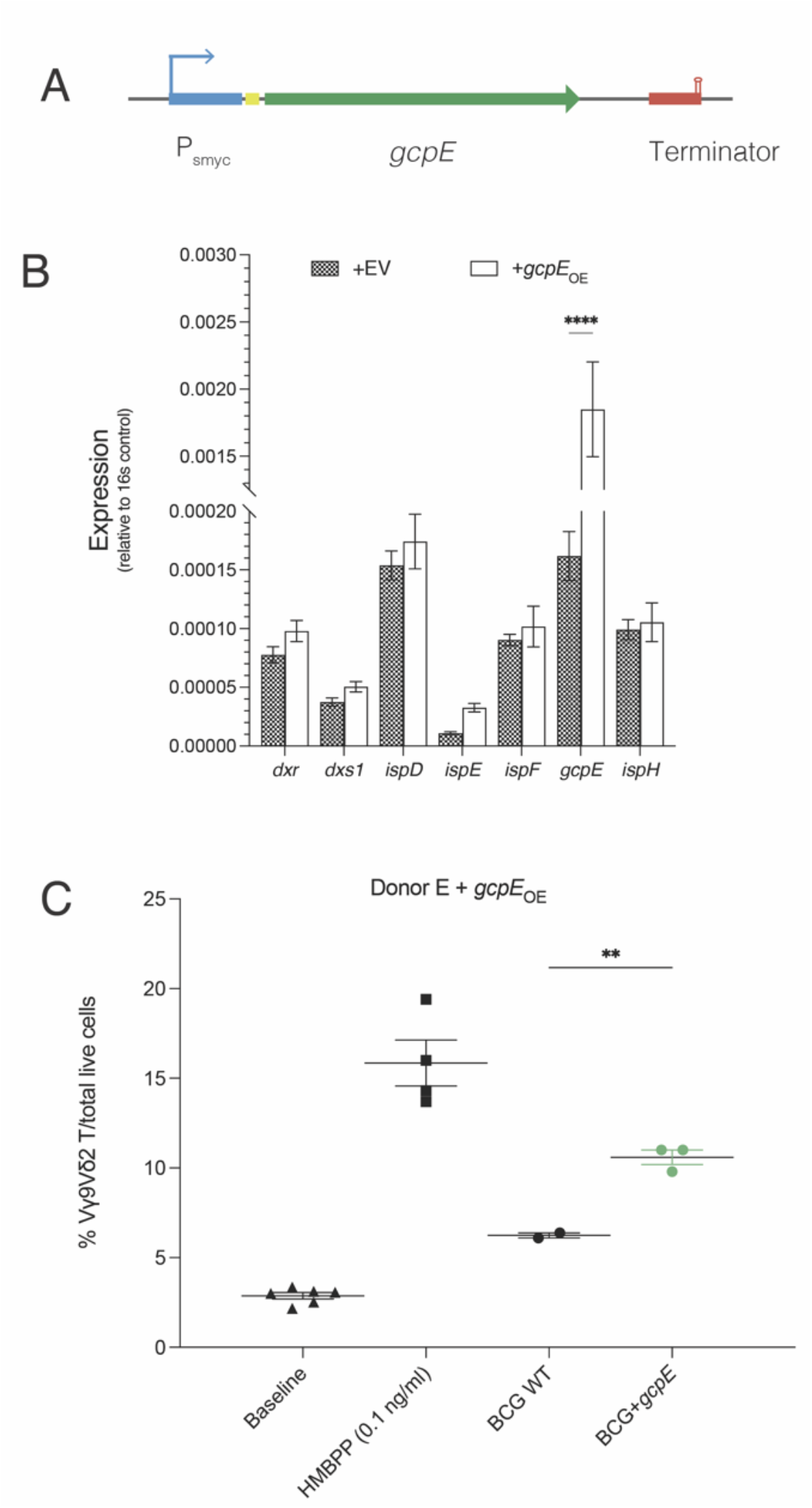
Generation of a *gcpE* overexpression construct. **A. Designing a *gcpE* OE construct**. The pCQ105 construct contains a single-gene expression platform composed of a strong mycobacterial promoter (blue) driving the expression of *gcpE* (green), which is preceded by a ribosome binding site (yellow) and followed by a terminator (red). **B. HMBPP synthase GcpE is strongly overexpressed**. Expression of each gene in the MEP pathway was compared between WT and pCQ105-expressing BCG. Compared to WT, *gcpE* was significantly overexpressed in the engineered strain (Sidak’s multiple comparisons; p<0.0001****). N=3 biological replicates in technical triplicate. **C. Overexpression of *gcpE* significantly enhances Vγ9Vδ2 T cell expansion over WT**. Shown is percent expansion of Vγ9Vδ2 T cells by pure HMBPP, 1:500 diluted lysate from either WT or BCG+pCQ105, compared to baseline. There is a significant difference in Vγ9Vδ2 between WT and BCG+pCQ105 (Sidak’s multiple comparisons; p<0.0001**). Shown is one PBMC donor, representative of independent experiments across two unique donors.

We transformed the plasmid pCQ105 carrying this construct into BCG and the resultant strain had no apparent growth defect. Using our *ex vivo* Vγ9Vδ2 T cell stimulation assay, we tested this strain and found that it enhanced activation of Vγ9Vδ2 T cells compared to WT (Figure 3C). This is in agreement with findings in *Mtb*, in which *gcpE* overexpression strongly enhanced activation over WT *Mtb* (34). Together, these data suggest that a targeted overexpression strategy may circumvent the tight regulation of this pathway in BCG. Importantly, it is critical to the success of BCG engineering to determine how to overcome broader feedback inhibition to maximize HMBPP production, and thus vaccine immunogenicity, in this system.

## Discussion

HMBPP is an intermediate of isoprenoid metabolism via the MEP pathway and potently activates and expands host Vγ9Vδ2 T cells (Fig 1). We found that this metabolic pathway is essential in BCG (Fig 1B), as has been reported in *Mtb* (34, 61, 62). Because Vγ9Vδ2 T cells are protective in the context of *Mtb* infection, we sought to generate a BCG vaccine strain that induced a stronger Vγ9Vδ2 response by synthesizing a bolus of HMBPP. Synteny analyses across mycobacterial genomes revealed certain pair biases of MEP genes, which we used to assemble a synthetic MEP locus (Fig 2A), in addition to a single gene overexpression construct (Fig 3A). We found that expressing the synthetic MEP locus caused feedback inhibition of the pathway (Fig 2B) and did not enhance Vγ9Vδ2 T cell expansion over the wild-type vaccine strain (Fig 2C, D). In contrast, the *gcpE* overexpression strain significantly expanded Vγ9Vδ2 T cells over WT BCG with no concomitant downregulation of other MEP genes (Fig 3B,C). Interestingly, we detected overexpression of the kinase IspE in both engineered strains (Figures 2B, 3B), presenting a potential role of this gene in the regulation of these pathways. Further improvements can be made to this strain through the addition of D-glyceraldehyde-3-phosphate, which has been shown in other systems to strongly accelerate Dxs activity (68).

Our engineering strategy is distinct from other BCG-based approaches. We modulated the bacterium’s isoprenoid biosynthesis pathway to accumulate a metabolic intermediate, which has not been attempted before in this system. While other approaches focus on antigen presentation, our work targets a metabolic, self-nonself sensing pathway to modulate immunity. Our initial attempts, including the multigene locus and *dxr* overexpression, disrupted the delicate balance of this pathway and presented barriers to engineering. Thus, understanding the molecular underpinnings of this phenomenon might reveal mechanisms by which we can manipulate the system to accumulate more HMBPP. Together, this study demonstrates the essentiality of the MEP pathway and underscores the importance of carefully-executed engineering of isoprenoid biosynthesis in BCG.

Future work is needed to address the metabolic flux of this pathway in the engineered strains, including metabolomic analysis of pathway intermediates, including HMBPP. Further, we will leverage a live infection model to determine whether the intracellular concentration of HMBPP, determined in this study via Vγ9Vδ2 T cell expansion, is relevant in the context of vaccination. It also remains unclear how HMBPP is released from phagocytosed bacteria into the cytosol of the macrophage. The bacterium might be actively secreting HMBPP, although this is unlikely because it is a polar, charged molecule and no transporter has been identified. Alternatively HMBPP could be released through bacterial lysis in the phagosome. It also remains to be determined whether HMBPP sensing requires access of the bacterium to the host cell cytosol via phagosomal perforation, or whether there is a yet-undetermined mechanism for HMBPP itself to cross the phagosomal membrane.

Finally, in order to optimize future engineering efforts using this strategy, it is critical to understand the mechanisms of isoprenoid feedback regulation and, importantly, how to overcome them. While much remains to be done on the path to developing a more protective vaccine against tuberculosis disease, we demonstrated the essentiality of the MEP pathway and explored potential avenues for metabolic reconstruction of BCG. Together, this study provides a novel investigation of isoprenoid biosynthesis pathway regulation in BCG, laying the groundwork to inform a future, rationally-reengineered vaccine strain.

## Acknowledgements

Many thanks to Preethi Thattai Ragunathan for her expertise and troubleshooting assistance, as well as Anne Xu, Miles Wingfield, Sarah Stanley, Russell Vance, Aziz Qabar, Astrid, and all members of the Cox lab for their helpful discussions and feedback. This work was supported by the National Institutes of Health (NIH T32GM132022) and the Henry Wheeler Center for Emerging and Neglected Diseases (CEND).

## Figure legends

**Supplementary Figure 1.**
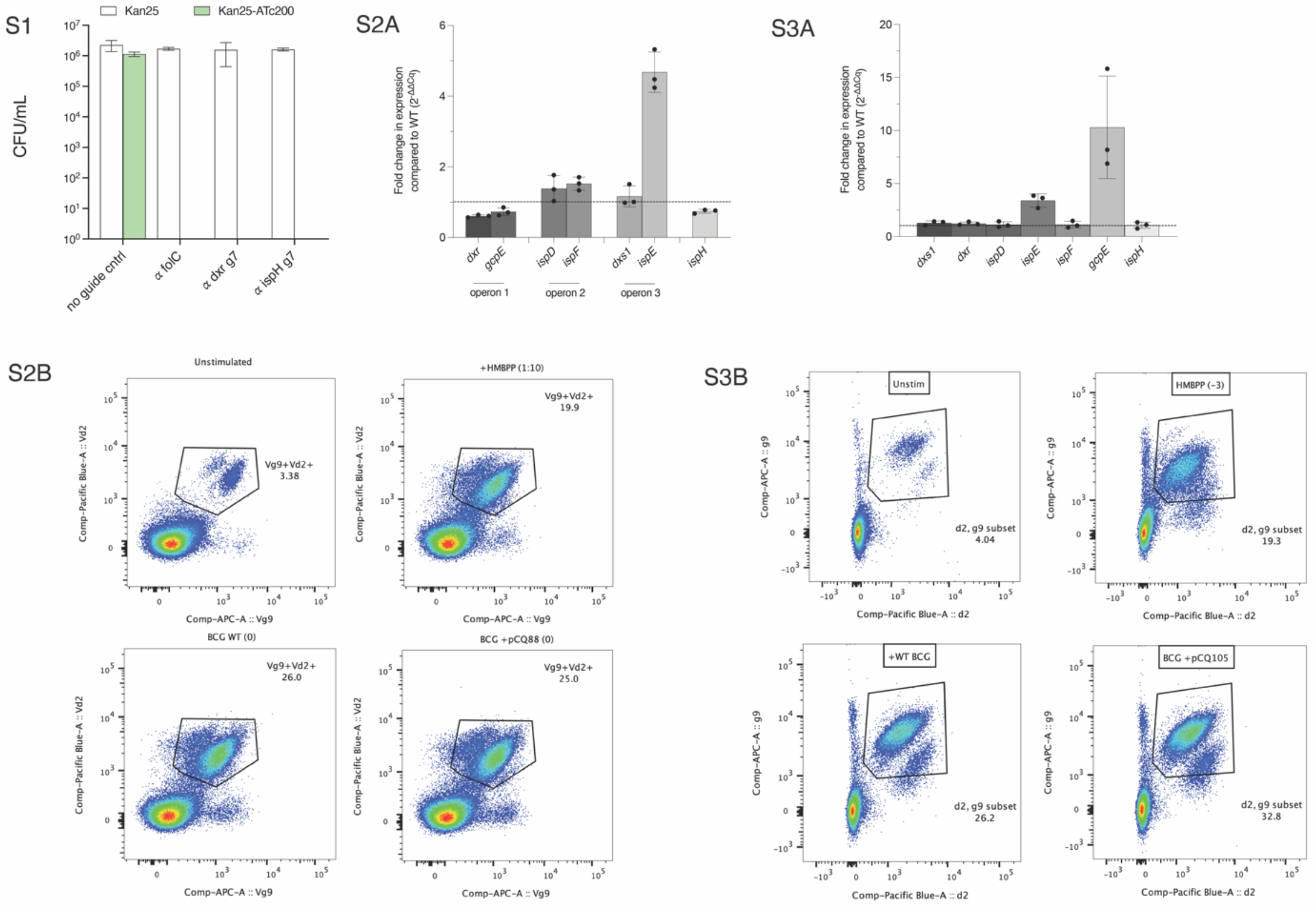
Quantification of CRISPRi targeting MEP in BCG. MEP genes *dxr* and *ispH* were transcriptionally repressed using inducible CRISPRi. CFU from 3 biological replicates, both uninduced and induced, were counted and quantified. Data plotted are mean CFU ± SEM.

**Supplementary Figure 2**

**A. Expression of MEP genes in synthetic locus relative to WT.** No significant differences in relative expression exist between WT and BCG+pCQ88. N=3 biological replicates in technical triplicate.

**B. Representative flow cytometry plots**. Shown are representative scatter plots of untreated, HMBPP-treated, and either WT BCG or BCG+pCQ88 (MEP_synth_) lysate-treated PBMCs. Cells were gated as follows: lymphocytes > singles > live > CD3+ > Vγ9/Vδ2 +/+. Shown are the results of the final gating of Vγ9+/Vδ2+ double-positive cells. All dot plots shown are from donor E and representative of 3 independent PBMC donors.

**Supplementary Figure 3**

**A. Expression of *gcpE* relative to WT.** Compared to WT, *gcpE* was strongly overexpressed in the engineered strain. N=3 biological replicates in technical triplicate. **B. Representative flow cytometry plots**. Shown are representative scatter plots of untreated, HMBPP-treated, and either WT BCG or BCG+pCQ105 (*gcpE*) lysate-treated PBMCs. Cells were gated as follows: lymphocytes > singles > live > CD3+ > Vγ9/Vδ2 +/+. Shown is the final gating of Vγ9+/Vδ2+ double-positive cells. All dot plots shown are from donor E and representative of 2 unique PBMC donors.

